# Untrimmed ITS2 metabarcode sequences cause artificially reduced abundances of specific fungal taxa

**DOI:** 10.1101/2024.08.02.606430

**Authors:** Kathleen E. Kyle, Jonathan L. Klassen

## Abstract

Advances in DNA metabarcoding have greatly expanded our knowledge of microbial communities in recent years. Pipelines and parameters have been tested extensively for bacterial metabarcoding using the 16S rRNA gene and best practices are largely established. For fungal metabarcoding using the ITS gene, however, only a few studies have considered how such pipelines and parameters can affect community prediction. Here we report a novel bias uncovered during ITS2 sequencing of *Trichoderma*-infected ant fungus gardens and confirmed using mock communities. Abnormally low forward read quality caused *Trichoderma* ITS2 reads to be computationally filtered before and during read pair merging, thus almost entirely eliminating *Trichoderma* ASVs from the resulting fungal community profiles. Sliding window quality trimming before filtering allowed most of these reads to pass filtering and merge successfully, producing community profiles that now correlated with visual signs of *Trichoderma* infection and matched the composition of the mock communities. Applying such sliding window trimming to a previously generated environmental ITS2 dataset increased the detected fungal diversity and again overcame read quality biases against *Trichoderma* to instead detect it in nearly every sample and often at high relative abundances. This analysis additionally identified a similar, but distinct, bias against a second fungal genus *Meyerozyma*. The prevalence of such quality biases against other fungal ITS sequences is unknown but may be widespread. We therefore advocate for routine use of sliding window quality trimming as a best practice in ITS2 metabarcoding analysis.

**Importance:** Metabarcode sequencing produces DNA abundance profiles that are presumed to reflect the actual microbial composition of the samples that they analyze. However, this assumption is not always tested, and taxon-specific biases are often not apparent, especially for low-abundance taxa in complex communities. Here we identified ITS2 read quality aberrations that caused dramatic reductions in the relative abundances of specific taxa in multiple datasets characterizing ant fungus gardens. Such taxon-specific biases in read quality may be widespread in other environments and for other fungal taxa, thereby causing incorrect descriptions of these mycobiomes.

## Introduction

Fungal classification is notoriously difficult (1, 2), which may partly explain why fungi remain understudied compared to bacteria (3–6) despite their global importance in terrestrial and plant-associated ecosystems, including agriculture (7–9). Community amplicon sequencing, or “metabarcoding”, has been widely used to characterize bacterial communities using the 16S rRNA gene as a common bacterial barcode (10–12). Metabarcoding was later adapted to fungal communities, especially using the internal transcribed spacer (ITS) region of the eukaryotic rRNA gene cluster (13). Despite its later adoption, ITS metabarcoding is now one of the most widely used techniques for characterizing microfungal communities (14–16).

There are many biases associated with DNA metabarcoding (17–19). From the method of DNA extraction to the many challenges of PCR, biases in library generation and sequencing are well-documented (14, 17, 20–24). Historically, most studies of computational biases focused on the algorithms used to bin sequences into operational taxonomic units (OTUs) or exact/amplicon sequence variants (E/ASVs) (25–27), and those used for taxonomic classification (28, 29). Some recent studies have also considered biases associated with the upstream data manipulation steps (30–34). Performed before the more computationally intensive binning and classification steps, such “preprocessing” checks the raw sequencing data for quality concerns and, if overlapping paired-end reads are available, merges reads into a consensus sequence.

Preprocessing standards have largely been developed for and adopted from bacterial metabarcoding (35–39). However, fungal ITS sequences pose extra challenges, particularly due to their length heterogeneity. For example, the widely used v4 region of the bacterial 16S rRNA gene (hereafter “16S v4”) is consistently ∼250 bp long, but the comparable fungal rRNA internal transcribed spacer region 2 (hereafter “ITS2”) can range from ∼50-800 bp long (40, 41). Thus, Illumina paired-end reads used to sequence 16S v4 metabarcodes overlap significantly with each other such that once merged, nearly every base is sequenced twice for improved basecall accuracy. When using a similar sequencing approach for ITS2 metabarcoding, sequences ∼250 bp long can be merged and therefore sequenced twice. However, if the ITS2 gene is < 250 bp the sequencing reads will contain non-biological bases when sequencing progresses into the 3’ adapter/primer region (“primer readthrough”; 42) and possibly beyond. In contrast, if the ITS2 sequences are too long the paired reads will share little or no overlap, reducing the proportion of bases sequenced twice and making merging difficult or impossible (43). This has led some researchers to question the value of using paired-end sequencing of ITS2 barcodes (44–46).

Successful read merging also depends on read quality. During preprocessing, low-quality bases are typically “trimmed” from the beginning and end of each read and primer readthrough is sometimes “clipped”. Entire reads that still don’t meet specified quality thresholds are then “filtered” out from the dataset. Because the length of the 16S v4 metabarcode is homogeneous, many pipelines remove the same number of bases from the ends of all reads to remove bases at the 3’ read end that have the lowest quality scores, producing higher quality reads of a fixed length (36, 37, 47–51). In contrast, truncation of ITS2 reads to a fixed length is inappropriate because they have more variable lengths. Thus, ITS pipelines often omit this truncation step (although some do clip primer readthrough; 42, 46, 52, 53).

Using untrimmed metabarcoding reads has several downsides. Filtering that only considers the average quality of entire reads will often remove many untrimmed reads that typically have lower-quality bases at their 3’ ends, notwithstanding many high-quality bases at their 5’ ends. Untrimmed reads that pass filtering will be more challenging to merge properly, if at all, due to the inclusion of these low-quality bases and adapter and primer sequences if they were not clipped. Any merged sequences will inherit the lower quality of these untrimmed reads, potentially leading to inaccurate taxonomic binning and classification. Filtering ITS2 metabarcode data may be especially inaccurate if pipeline parameters are unchanged from defaults chosen for 16S v4 datasets. The presence of low-quality data may also increase compute times. More fundamentally, using uniform read lengths assumes that read quality varies similarly for all reads in a dataset. This assumption seems weak given the taxon-specific biases of most other metabarcoding steps (12, 14, 54–56).

Here we report an underappreciated taxon-specific filtering bias during ITS2 metabarcoding (but see ref #57). In our previous research (58), we infected ant fungus gardens with *Trichoderma* (Ascomycota: Sordariomycetes: Hypocreales: Hypocreaceae), leading *to Trichoderma* growth that was visually apparent, yet there were no *Trichoderma* reads after ITS2 metabarcoding. We determined that *Trichoderma* reads in these samples uniquely failed to pass the filtering and read merging steps in our analysis pipeline, removing nearly all *Trichoderma* sequences from the final output. Using sliding window quality trimming before filtering remedied this bias against *Trichoderma* in both defined mock communities and our experimental infection dataset. Furthermore, we detected the same bias against *Trichoderma* in an ITS2 metabarcoding dataset from environmental samples, as well as a similar bias against *Meyerozyma* (Ascomycota: Saccharomycetes: Saccharomycetales: Saccharomycetaceae) that was also remedied by sliding window quality trimming. This study demonstrates how a taxon-specific bias due to an unusual reduction in quality at the 3’ end of ITS2 metabarcoding reads was not accommodated by typical filtering parameters, which led to erroneous taxonomic profiles and thus erroneous biological conclusions.

## Results

During our previous work infecting fungus gardens cultivated by *Trachymyrmex septentrionalis* ants with the fungal pathogen *Trichoderma* (58), we visually observed *Trichoderma* growth on infected fungus gardens and not on control fungus gardens inoculated only with buffer (Fig. 1A). We were therefore surprised when our ITS2 community profiles for the infected fungus gardens contained very few *Trichoderma* ASVs (Fig. 1B, Suppl. Fig. S1A), despite their containing the expected ASVs from the ant’s cultivar fungus (the main constituent of ant fungus gardens). Compared to the mock-inoculated controls there were also many fewer reads in infected samples after filtering, with nearly all reads removed after paired read merging (Fig. 1C, Suppl. Fig. S1B). The infected samples also had distinctively abnormal read quality profiles. At ∼125 bp, forward read quality suddenly became highly variable and the median decreased sharply from a Phred score of ∼35 to a score of ∼25 that then persisted to the ends of these 250 bp reads (Fig. 1D, Suppl. Fig. S1C). The quality of the reverse reads was poorer than the quality of the forward reads in every sample, but the quality of reverse reads from infected samples was noticeably poorer than that of those from the uninfected samples (Suppl. Fig. S1C). These data suggested an unexpected and dramatic bias against ITS2 reads from *Trichoderma* in this analysis.

**Figure 1.**
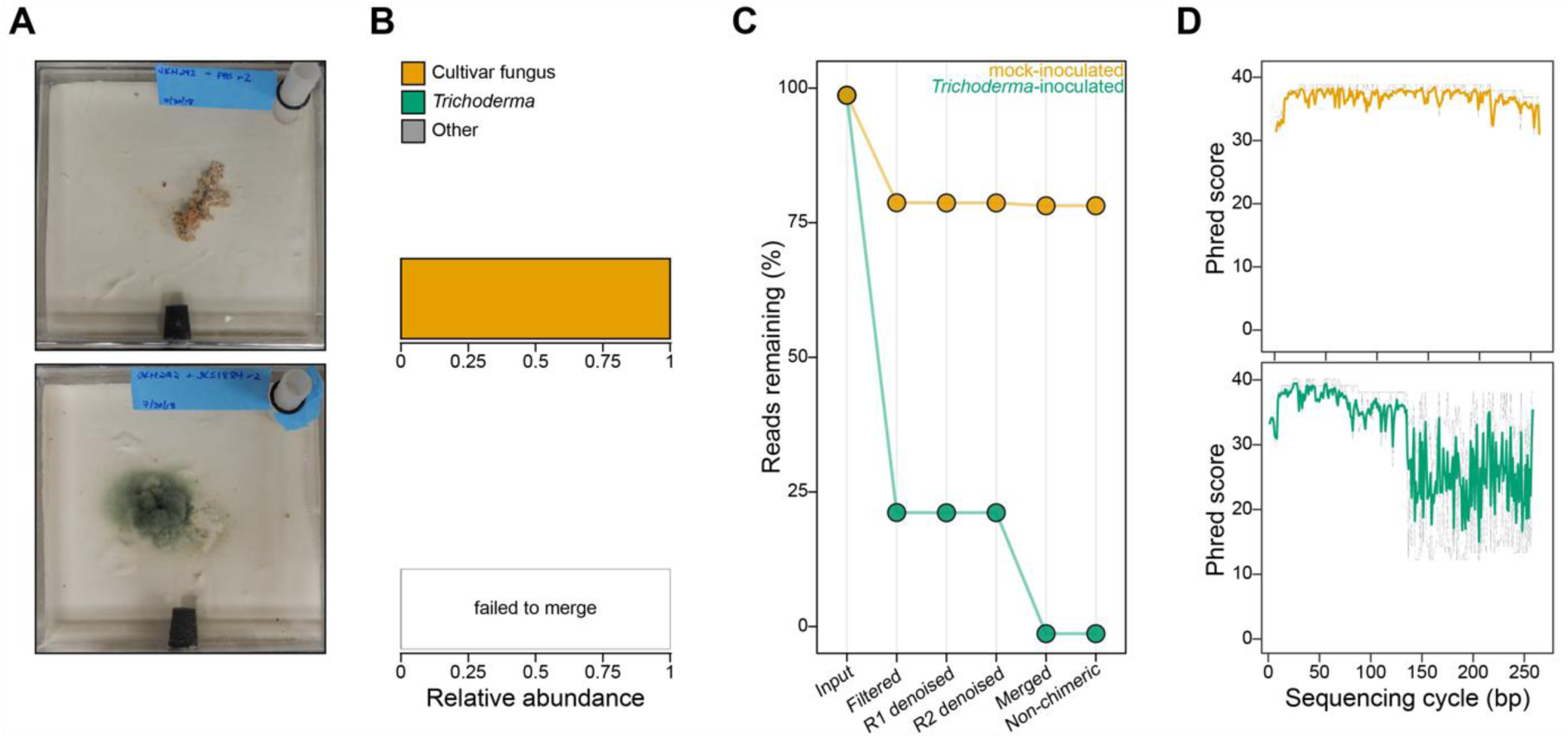
*Trichoderma* ITS2 reads are completely removed by filtering or failure to merge during metabarcode sequencing analysis and have abnormal forward read quality. **A)** Representative images of a healthy fungus garden 4 days after inoculation with sterile PBS (“mock-inoculated”, top) and an infected fungus garden 4 days after inoculation with *Trichoderma* spores in PBS (“*Trichoderma*-inoculated”, bottom). **B)** Relative abundances of fungal ASVs for the mock-inoculated (top) and *Trichoderma*-inoculated (bottom) fungus gardens pictured in A. **C)** Percent of reads remaining after each step of the metabarcoding analysis pipeline for the samples pictured in A. Forward and reverse reads are abbreviated as R1 and R2, respectively. **D)** Quality plots for forward ITS2 reads from the samples pictured in A. Reads from the mock-inoculated (top) and *Trichoderma*-inoculated (bottom) samples are plotted in orange and green, respectively. The solid line shows the median basecall quality score (Phred) at each base position and dotted gray lines show the basecall quality quartiles. See Suppl. Fig. S1 for results from the full dataset.

To reproduce this *Trichoderma*-specific bias more quantitatively, we sequenced mock communities created using defined proportions of DNA from pure cultures of *Trichoderma* and the ant cultivar fungus. As expected, the reduction in quality midway through the forward reads and the number of reads discarded during filtering and merging both increased alongside the proportion of *Trichoderma* DNA in the mock communities (Fig. 2A, Suppl. Fig. S2). No *Trichoderma* ASVs were detected in any of the mock communities, including that containing 100% *Trichoderma* DNA (Fig. 2B). Somewhat surprisingly, we detected cultivar fungus ASVs in the 100% *Trichoderma* mock community, but at very low levels likely originating from cross-contamination during community generation or sequencing.

**Figure 2.**
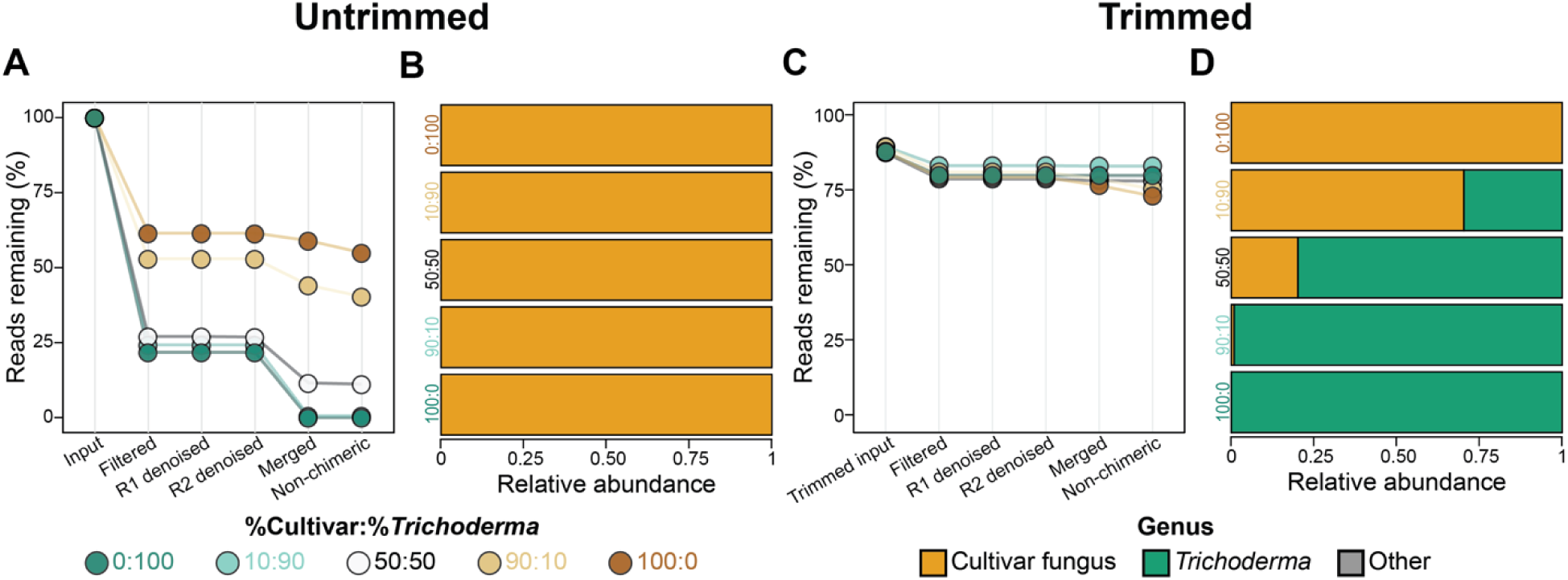
Mock communities replicate the bias against *Trichoderma* ITS2 metabarcodes and demonstrate that sliding window trimming can mitigate it. **A)** Percent of untrimmed reads remaining at each step of the metabarcoding analysis pipeline for mock communities constructed using different proportions of cultivar fungus and *Trichoderma* DNA. **B**) ASV relative abundances in each mock community using untrimmed reads. **C**) Percent of trimmed reads remaining at each step of the metabarcoding analysis pipeline for the same mock communities as in (A). **D**) ASV relative abundances in each mock community using trimmed reads. Forward and reverse reads are abbreviated as R1 and R2, respectively.

We hypothesized that the sudden quality drop in the middle of the forward *Trichoderma* reads caused most of them to fall below the default quality thresholds and thus be filtered out or fail to merge, causing low final *Trichoderma* read counts and the absence of *Trichoderma* ASVs. To test this, we first quality trimmed the 3’ end of all reads in the mock community samples using Trimmomatic’s sliding window trimmer (59) and then analyzed them as before. Sliding window quality trimming reduced the drop in read quality midway through the forward reads (Suppl. Fig. S3), and most trimmed reads successfully passed the read filtering and merging steps of the analysis pipeline (Fig. 2C). *Trichoderma* ASVs were now detected in all mock communities except for that containing 100% cultivar fungus DNA and their relative abundances correlated with the expected cultivar fungus:*Trichoderma* ratios of the input DNA (Fig. 2D), albeit with some overrepresentation of *Trichoderma*, perhaps due to different ITS2 copy numbers in *Trichoderma* and the cultivar fungus, not constructing the mock communities as molar proportions, or sequencing bias favoring *Trichoderma*. Overall, sliding window trimming generated metabarcoding community profiles that were much closer to the expected values compared to the profiles generated without such trimming (Fig. 2).

We next applied sliding window trimming to our initial *Trichoderma*-infection dataset. Now *Trichoderma* ASVs were detected at relative abundances consistent with the visual appearance of these samples (Suppl. Fig. S4A), and the number of trimmed reads that were retained after filtering and merging (Suppl. Fig. S4B) and the average quality of the trimmed forward reads (Suppl. Fig. S4C) both increased for all *Trichoderma*-infected samples. Thus, sliding window trimming successfully mitigated bias against *Trichoderma* in these experimental samples, as it did for the mock communities.

Finally, we tested the effect of sliding window trimming on an environmental ITS2 dataset that we previously generated from 98 freshly excavated ant fungus garden samples (58). Without trimming, very little taxonomic diversity appeared in this environmental dataset (Fig. 3, left), with 86/90 samples having ≥ 98% cultivar reads, 3 samples containing < 30% cultivar reads, and 1 sample having 62% cultivar reads. In contrast, considerably greater taxonomic diversity was apparent after sliding window trimming, particularly for *Trichoderma,* which occurred in nearly all trimmed samples (81 out of 90) at often high relative abundances (Fig. 3, right). Similarly, trimming increased the relative abundance of the yeast genus *Meyerozyma* in four samples. One sample had 19% *Meyerozyma* before trimming that increased to 88% after trimming, and three samples went from 0% to 1.5%, 67%, or 93% *Meyerozyma*. Log2-transformed fold changes of trimmed versus untrimmed relative abundances of the genera in these environmental samples confirmed that *Trichoderma* and *Meyerozyma* relative abundances increased following sliding window trimming (Fig. 4). In this analysis, the genera whose relative abundances decreased after read trimming (cultivar fungus, *Penicillium*, *Cladosporium*, and *Oberwinklerozyma*) occurred in samples also containing *Trichoderma*, *Meyerozyma*, and/or “other” fungi due to the relative nature of the metabarcoding data (i.e., the trimmed samples with the largest taxon increases also had the largest taxon decreases).

**Figure 3.**
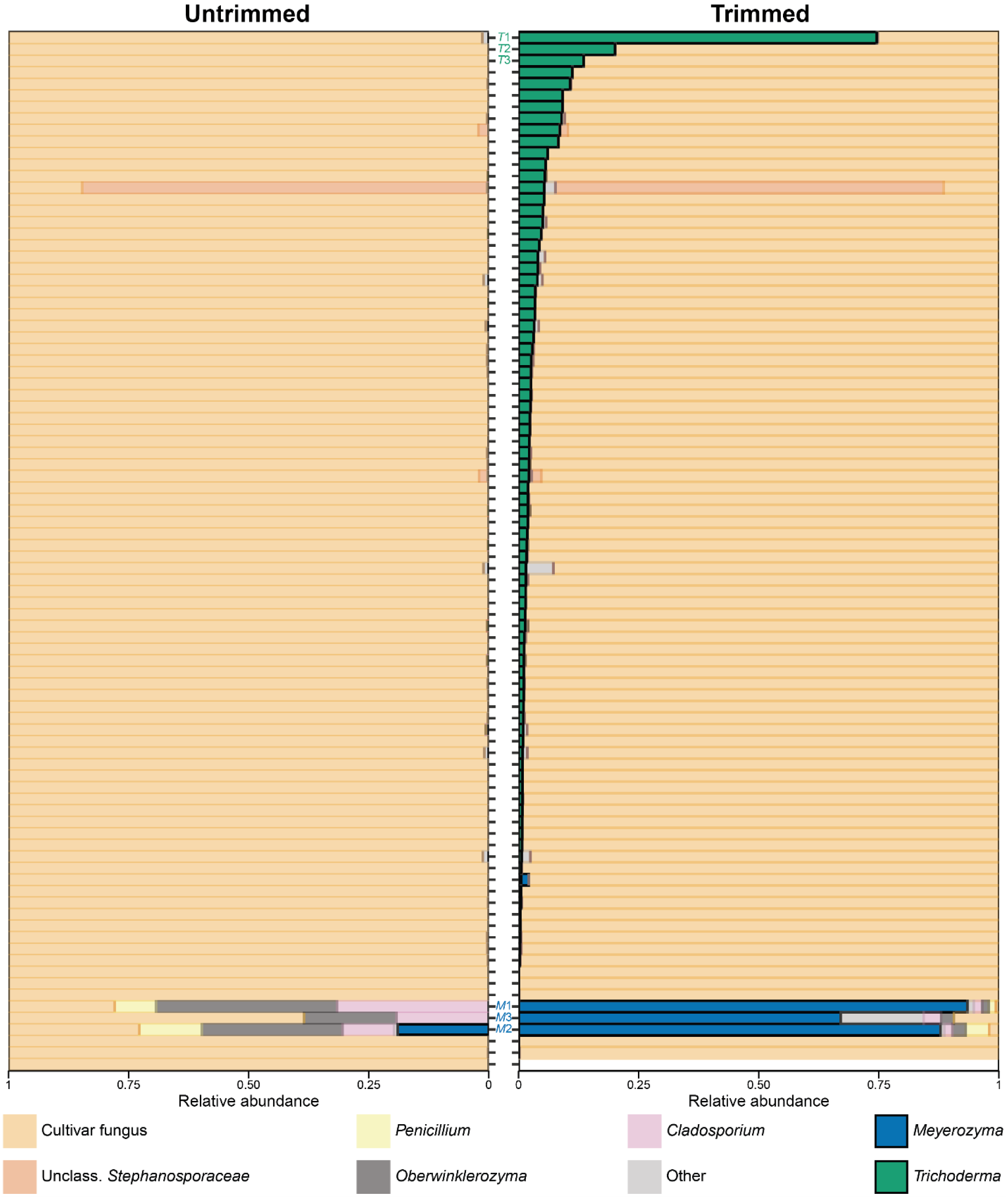
Using trimmed reads revealed greater diversity of non-cultivar fungi in environmental fungus gardens compared to using untrimmed reads. ASV relative abundances in environmental fungus gardens using untrimmed (left) and sliding window-trimmed (right) reads. Each row represents an individual fungus garden, which are ordered by the relative abundance of *Trichoderma* after trimming. The samples with the three highest relative abundances of *Trichoderma* or *Meyerozyma* are labeled *T*1-*T*3 and *M*1-*M*3, respectively. The taxon labelled “Other” includes all ASVs that were < 1% abundant in all samples. See Suppl. Fig. S5 for the absolute abundances of fungal reads in this dataset.

**Figure 4.**
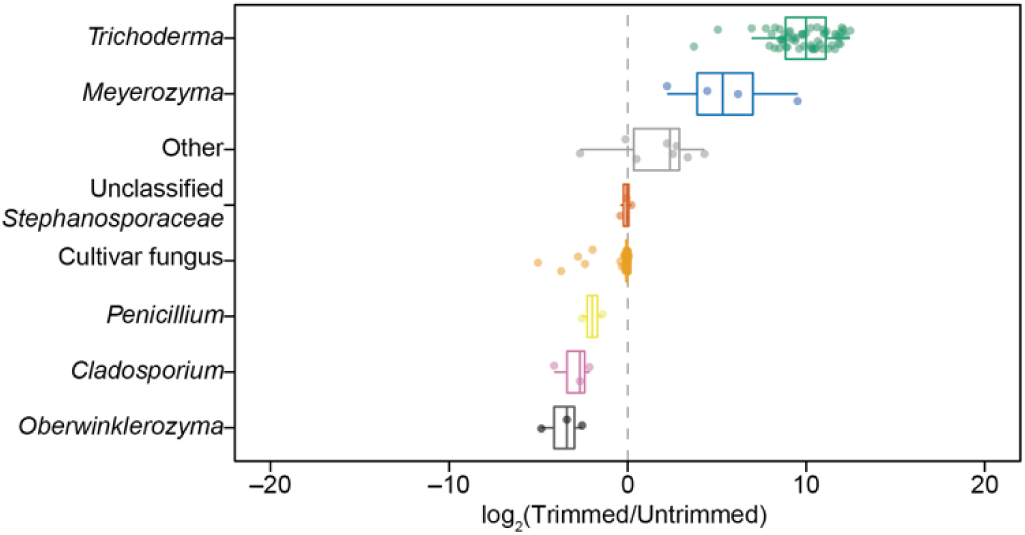
Trimming particularly increases the relative abundances of *Trichoderma* and *Meyerozyma*. The relative abundances of genera in the environmental samples (Fig. 3) were compared using their Log2-transformed fold-change in trimmed versus untrimmed datasets, with each dot comparing the relative abundance of a genus in a single fungus garden sample. The center line and outer edges of the boxplots show the median and quartiles of the fold-change in relative abundances for each genus, respectively. Whiskers extend to the highest and lowest data points no further than 1.5x the interquartile range. Genera were excluded if they were < 1% abundant in both untrimmed and trimmed datasets. Taxon labels and colors match those in Fig. 3.

Closer inspection of the environmental samples with the highest abundances of either *Trichoderma* (*T*1-*T*3) or *Meyerozyma* (*M*1-*M*3) mirrored the changes we observed in our infection and mock community experiments following sliding window trimming (Fig. 5). Left untrimmed, these environmental samples all had distinctive drops in the quality of the forward reads (Fig. 5B, G) and high numbers of reads discarded during filtering and merging (Fig. 5C, H), making both taxa underreported in the relative abundance plots (Fig. 5E, J). Notably, the quality plots for samples containing *Meyerozyma* were similar to, but distinct from, those containing *Trichoderma,* with the sudden quality drop of the forward reads occurring at ∼200 bp for *Meyerozyma* (Fig. 5G) compared to at ∼125 bp for *Trichoderma* (Fig. 5B, Fig. 1C, Suppl. Fig. S1C). Sliding window trimming mitigated both taxon-specific biases by improving the quality of the forward reads (Fig. 5A, F) and increasing read retention during read filtering and merging (Fig. 5C, H). The resulting relative abundances of both taxa were much higher (Fig. 5D, I) compared to in the untrimmed samples. Given our detection of such biases in samples that contained only limited taxonomic diversity, we speculate that similar read quality biases may be widespread in metabarcoding datasets that include many other fungal taxa.

**Figure 5.**
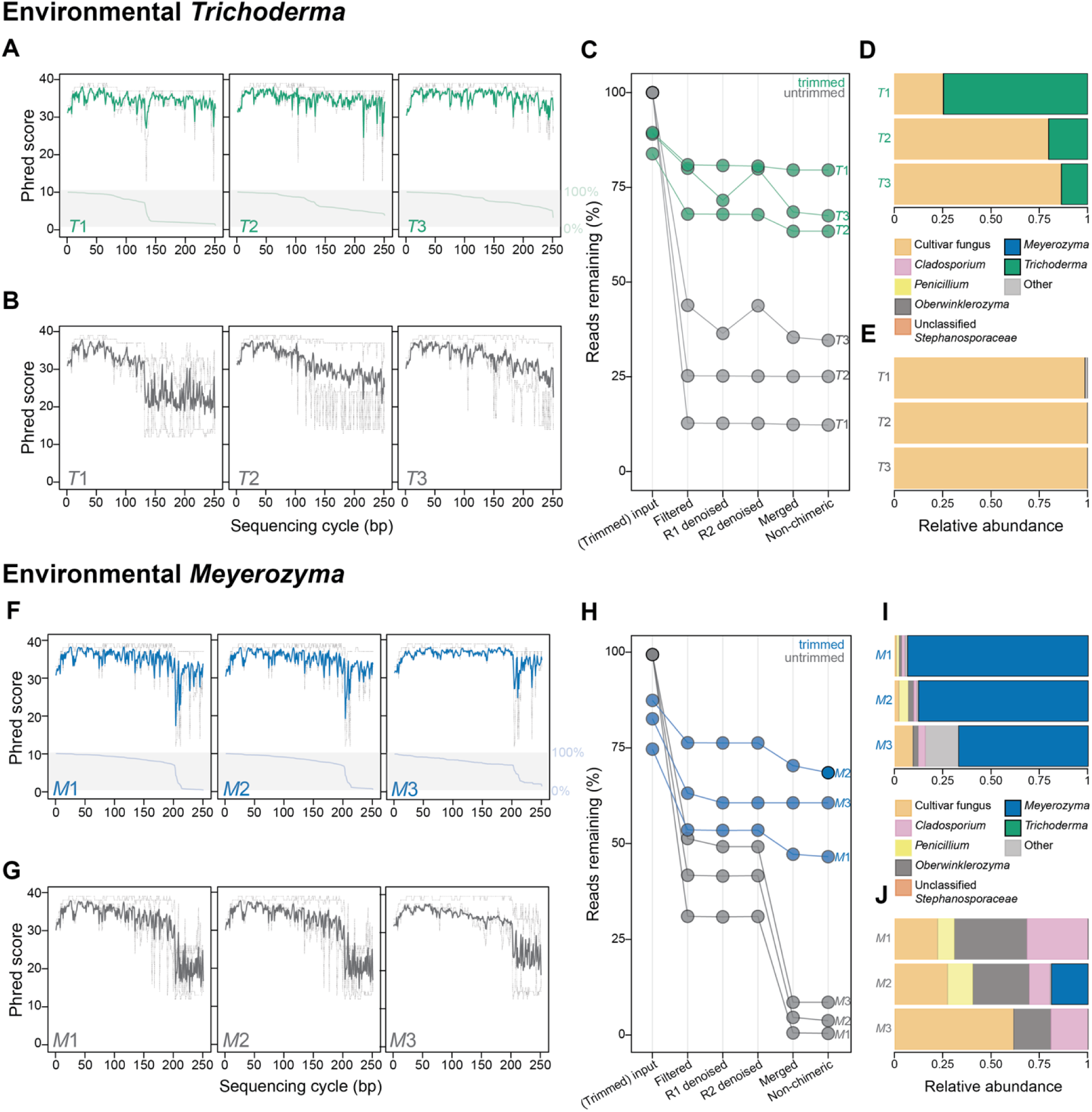
Trimming increases forward read quality and the number of reads that pass filtering and merge successfully for the environmental samples with the highest relative abundances of *Trichoderma* (*T*1-*T*3) and *Meyerozyma* (*M*1-*M*3). **A/B)** Forward read quality plots for trimmed (A) and untrimmed (B) samples *T*1-*T*3. **C)** Percent of reads remaining after each analysis pipeline step for trimmed and untrimmed samples *T*1-*T*3. **D/E)** Relative abundances of fungal ASVs in trimmed (D) and untrimmed (E) samples *T*1-*T*3. **F/G)** Forward read quality plots for trimmed (F) and untrimmed (G) samples *M*1-*M*3. **H)** Percent of reads remaining after each analysis pipeline step for trimmed and untrimmed samples *M*1-*M*3. **I/J)** Relative abundances of fungal ASVs in trimmed (I) and untrimmed (J) samples *M*1-*M*3. In panels A, B, C, F, G, and H, untrimmed data are colored gray and trimmed data are colored green (samples *T*1-*T*3) or blue (samples *M*1-*M*3). In panels A, B, F, and G, the median quality and the quality quartiles per base are plotted as solid and dotted lines, respectively. The light green or blue lines at the bottom of panels B and F, respectively, show the percentage of reads trimmed to that sequence length or longer. Forward and reverse reads are abbreviated as R1 and R2, respectively.

## Discussion

This study demonstrates that data preprocessing, particularly sliding window quality trimming, can significantly affect fungal metabarcoding data quality and analytical output. The taxon-specific ITS2 read quality biases that we identified (Figs. 1D, 5B, 5G, Suppl. Figs. S1C, S2) required these reads to be trimmed without applying a fixed truncation length to the entire dataset to avoid taxon-specific read loss during downstream filtering and merging (Figs. 1C, 2A, 5C, 5H, Suppl. Fig. S1B) and ultimately the underrepresentation or absence of these taxa in the resulting ITS2 community profiles (Figs. 1B, 2B, 3, 5E, 5J, Suppl. Fig. S1A). In fact, sliding window quality trimming improved read retention after filtering and merging for all taxa in our datasets (Fig. 2C, Suppl. Figs. S4B, S5) and improved classification of the cultivar fungus (Suppl. Fig. S6), demonstrating its general benefit compared to fixed length truncation. So long as a metabarcoding pipeline can tolerate reads with variable lengths, sliding window quality trimming should apply to metabarcoding using any barcode. We did not detect “ASV splitting” following trimming (Suppl. Fig. S6), which has been thought to arise from variable metabarcode read lengths (60, 61). Concatenating paired reads, alongside or instead of merging, may also improve taxonomic classification (30, 60), particularly for long amplicons that do not overlap enough for merging (34). That most reads in our study successfully merged after trimming (Figs. 2C, 5C, 5H, Suppl. Fig. S4B) suggests that the abnormally low forward read quality we observed is instead likely due to sequencing progressing beyond the end of short ITS2 template DNA molecules.

Our results additionally emphasize the importance of sequencing mock community controls alongside experimental samples (Fig. 2). Mock community controls are a gold standard tool to identify taxon-specific biases and validate computational pipelines (62). Ideally, these communities should contain every taxon present in the experimental communities being analyzed. Paradoxically, this requires prior knowledge of experimental community composition that is often unavailable, especially for understudied communities. Alternatively, constructing mock communities that include all currently known taxa is as impractical as it is impossible. Custom mock communities offer some promise (30, 63, 64), but these still require *a priori* knowledge of community composition, are technically challenging to create, are not standardized across research groups, and are currently rare for fungi (ATCC MSA-1010 and MSA-2010 from American Type Cultivar Collection, Manassas, VA, USA; 40, 65). Even if not comprehensive, mock community controls are useful to detect biases against the taxa that they do contain and thus should be used routinely.

When representative mock community controls do not exist, mindful data analysis is imperative. Researchers analyzing metabarcoding data should perform sufficient quality control during all steps of a computational analysis and especially appreciate that the default parameters are often set using well-characterized bacterial communities that may not apply to studies of other communities, particularly those targeting fungi (see 57 for an example of parameter optimization). These quality checks should include, but are not limited to, evaluating each sample for unusual patterns of read quality, tracking how many reads are removed from samples at each pipeline step, and comparing final community composition metrics between datasets processed using different parameters (e.g., trimmed vs. untrimmed). Results also should be examined carefully with respect to prior expectations given the experimental design, for example during ITS2 metabarcoding studies attempting to detect *Trichoderma* in plant root communities following its application as a biocontrol agent (e.g., 66). Despite their limitations, metabarcoding studies of novel or under-characterized microbial communities are important and necessary.

Most importantly, this study demonstrates how a seemingly small adjustment to data preprocessing can significantly impact the biological conclusions drawn from an analytical interpretation. Without sliding window trimming, environmental *T*. *septentrionalis* fungus gardens appeared to have perplexingly little microfungal diversity (Fig. 3 left). In contrast, trimmed reads revealed that environmental fungus gardens host a more diverse and abundant fungal community, of which *Trichoderma* is exceptionally prevalent (Fig. 3 right, Fig. 4). This latter result better agrees with the presence of metabolites commonly associated with *Trichoderma* in ant fungus gardens (58), other studies that have cultured many different microfungi (including *Trichoderma*) from ant fungus gardens (67–73), and the known abundance of *Trichoderma* in diverse soil and plant-associated communities such as the rhizosphere (74–78) where it often acts as a mycoparasite (79–81). Unexpectedly, we further discovered a strikingly similar bias against ITS2 reads from an unrelated genus, *Meyerozyma* (Fig. 5F-J), that belongs to a different subphylum (Saccharomycotina) than *Trichoderma* (Pezizomycotina). In our other research projects, we found another similar quality bias against ITS2 reads from *Clonostachys* (Suppl. Fig. S7), a genus more closely related to *Trichoderma*, both in the order Hypocreales. Rolling et al. (57) reported a similar taxon-specific aberration in read quality for different taxa than those studied here and using ITS1 metabarcoding.

Therefore, although the full distribution of such biases across all fungal taxonomy and barcodes is unknown, these data suggest they could be widespread. In conclusion, we recommend using sliding window quality trimming, appropriate quality controls, and mindful data analysis as part of best practices for metabarcoding, particularly for fungi.

## Methods

### Data Generation

Except for the cultivar fungus:*Trichoderma* DNA mock communities, all data and methods have been described elsewhere (58). The mock communities were prepared by extracting genomic DNA from pure cultures of *Trichoderma* and the *T*. *septentrionalis* cultivar fungus isolated from laboratory colony JKH000219 which was collected from Florida in 2016 (Florida Department of Agriculture and Consumer Services unnumbered Letter of Authorization; 58). *Trichoderma* strain JKS001884 was grown on Potato Dextrose Agar (PDA, Difco) + antibiotics (ABX, 50 mg/L penicillin and 50 mg/L streptomycin; both Fisher Scientific) at 25 °C for 1 week. Cultivar fungus was isolated by collecting small tufts of hyphae from the JKH000219 fungus garden using sterile extra-fine forceps (being careful to only collect hyphae and not surrounding pieces of forage) and growing them on PDA+ABX plates at 25 °C. These were checked daily for pathogen (non-cultivar) growth, in which case pathogens were cut and removed from the agar plates using sterile blades or cultivar hyphae were transferred onto to new PDA+ABX plates. This continued until the cultivar fungus comprised a pure culture (∼2-4 weeks).

Hyphae from these pure cultures were collected into bead-beating tubes with 250 µL of cetyltrimethylammonium bromide (CTAB) buffer and 0.5 g each of 0.1 mm and 1 mm sterile silica/zirconium beads for DNA extraction, and extracted DNA was quantified using Qubit 3.0 with a dsDNA high sensitivity kit (58). Five mock communities were created using ratios of 100:0, 90:10, 50:50, 10:90, 0:100 cultivar fungus to *Trichoderma* genomic DNA by mass. These mock communities were submitted to the Microbial Analysis, Resources, and Services (MARS) facility at the University of Connecticut for ITS2 metabarcode sequencing using Illumina indexed primers fITS7 (aka ITS3, 5’-GTGARTCATCGAATCTTTG-3’, 63) and ITS4 (5’-TCCTCCGCTTATTGATATGC-3’, 82) that contained Illumina adapters and dual 8 base indices (49). Samples were amplified from 30 ng of extracted DNA in triplicate 15 µl reactions using Go-Taq DNA polymerase (Promega) with the addition of 3.3 μg bovine serum albumin (BSA, New England BioLabs). To overcome inhibition from host DNA, 0.1 pmol of each primer without the adapters or indexes was added to the mastermix. The ITS2 PCR reaction was incubated at 95 °C for 2 minutes, then for 5 cycles of 30 s at 95.0 °C, 60 s at 48.0 °C and 60 s at 72.0 °C, then for 25 cycles of 30 s at 95.0 °C, 60 s at 55.0 °C and 60 s at 72.0 °C, followed by final extension at 72.0°C for 10 minutes. PCR products were pooled for quantification and visualization using a QIAxcel with a DNA Fast Analysis cartridge (Qiagen). PCR products were normalized based on the concentration of DNA from 250-400 bp then pooled using the epMotion 3075 liquid handling robot. The pooled PCR products were cleaned using Omega Bio-Tek Mag-Bind Beads according to the manufacturer’s protocol using a ratio of 0.8x beads to PCR product. The cleaned pool was sequenced on the MiSeq using a v2 2×250 base pair kit (Illumina, Inc).

### Data Analysis

All ITS2 amplicon datasets were analyzed using R v3.6.3 or 4.1.0 (83). The untrimmed dataset was processed following the DADA2 “ITS Pipeline Workflow (1.8)” (37). The only changes were setting parameter “randomize” to “TRUE” for learning errors with function “learnErrors”, dereplicating the forward and reverse reads using function “derepFastq” before running the “dada” function, graphing the output of the “track” variable using R barplot to visualize read retention at each pipeline step, and using ITSx v1.1.3 (84) with default parameters to remove potential flanking 18S rRNA regions from the ASVs prior to taxonomic classification. ASVs were removed that did not pass ITSx filtering and duplicate ITSx-treated ASVs were merged. These ASVs were then classified as described in (37) using the UNITE database v8.2 general fasta format (85). The resulting ASV table, taxon table, and sample data table were collected into a phyloseq object for further processing using phyloseq v1.26.1 (37, 86) and tidyr v1.3.1 (87). For the trimmed dataset, reads were processed exactly as above except FASTQ files were first trimmed at the 3’ end using Trimmomatic v0.39 (59) with parameters SLIDINGWINDOW:5:20.

For each sample, ASVs that were < 1% abundant were manually classified as “other” and their abundances were combined. *T. septentrionalis* ants predominantly cultivate fungi from tribe *Leucocoprineae* (*Agaricaceae*: *Agaricales*), typically annotated as either genus *Leucocoprinus* or *Leucoagaricus* (88–90). However, the taxonomy of these fungi is complex (90–92) and thus ASVs were manually defined as “cultivar fungus” if they were classified as belonging to *Leucocoprinus*, *Leucoagaricus*, or “unclassified family *Agaricaceae*”. These cultivar fungus ASVs were confirmed to be closely related to other fungus-growing ant cultivar fungi using NCBI blastn megablast against the nonredundant nucleotide database, nr/nt (93, 94). All code used for analysis is available at https://github.com/kek12e/ms_ITS2trimming.

## Data availability

Unprocessed fungal ITS2 community amplicon sequencing FASTQ files are publicly available in the National Center for Biotechnology Information (NCBI) Sequence Read Archive (SRA) under BioProject PRJNA763335 (environmental fungus gardens), PRJNA743045 (*Trichoderma*-infected laboratory fungus gardens), and PRJNA1138067 (cultivar fungus:*Trichoderma* mock communities and *Clonostachys*).

## Acknowledgements

Funding for this work was received from NSF grant IOS-1656475 (J.L.K). We thank Dr. Kendra Maas from the Microbial Analysis, Resources, and Services (MARS) facility at the University of Connecticut for assistance with ITS2 library preparation and sequencing, and Madison Adams for generation of the environmental ITS2 dataset.

