## Supplemental Figures for "Untrimmed ITS2 metabarcode sequences cause artificially reduced abundances of specific fungal taxa"

### Supplementary Material

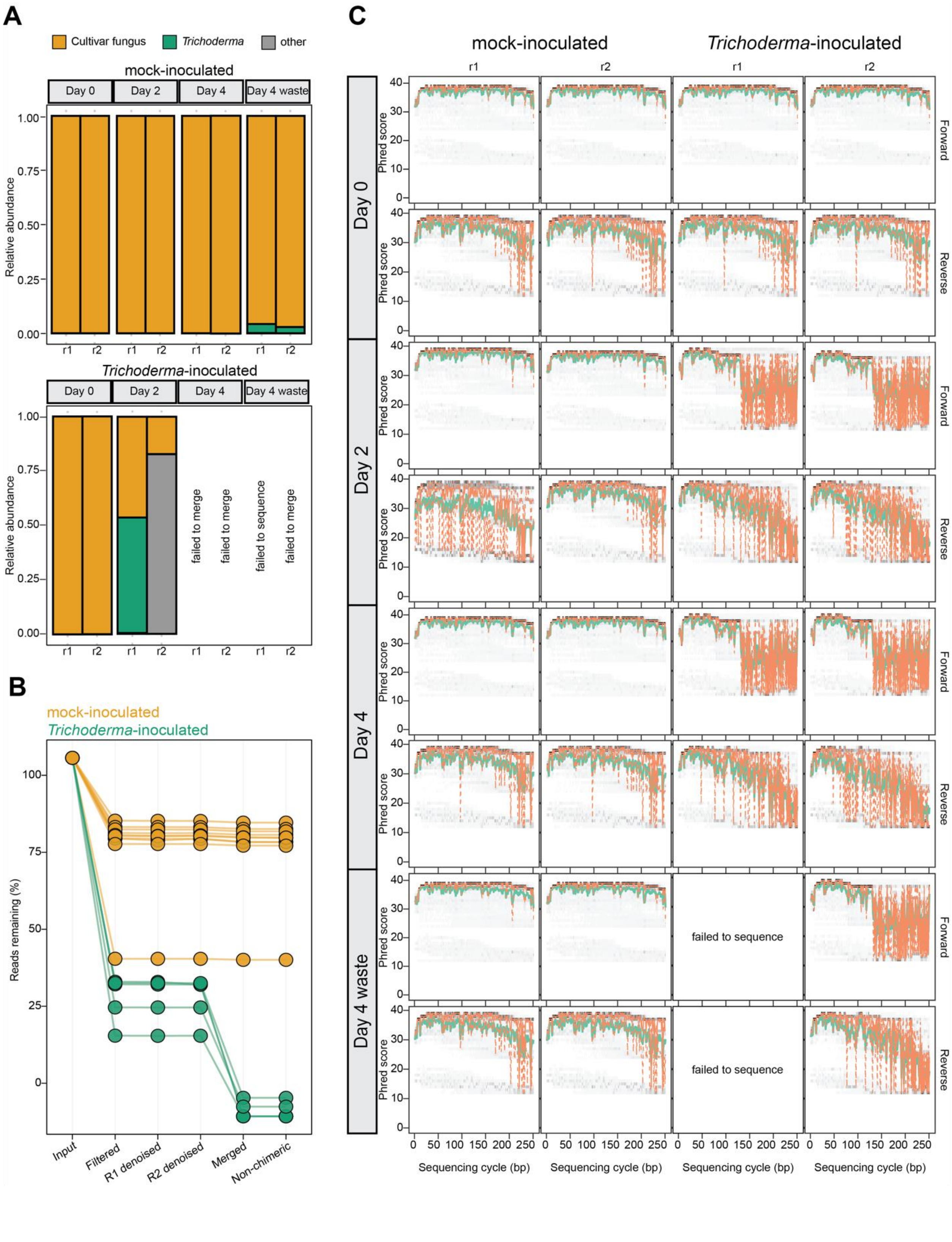

A
Cultivar fungus
Trichoderma
other
mock-inoculated
Day 0
Day 2
Day 4
Day 4 waste
Relative abundance
Trichoderma-inoculated
failed to merge
failed to merge
failed to sequence
failed to merge
B
mock-inoculated
Trichoderma-inoculated
Reads remaining (%)
Input
Filtered
R1 denoised
R2 denoised
Merged
Non-chimeric
C
mock-inoculated
Trichoderma-inoculated
Day 0
Day 2
Day 4
Day 4 waste
Forward
Reverse
Phred score
failed to sequence
failed to sequence
Sequencing cycle (bp)

**<u>Figure S1</u>. Untrimmed *Trichoderma* ITS2 reads are consistently filtered out or fail to merge due to unusually low read quality across the entire infection experiment. A)** Relative abundance stacked bar charts using untrimmed reads for the time course *Trichoderma* infection experiment shown in Fig. 1. Days 0, 2, and 4 follow inoculation with either *Trichoderma* spores (*“Trichoderma*-inoculated”) or sterile PBS (“mock-inoculated”). Fungus garden waste (i.e., pieces removed from the fungus garden by the ants) was only collected on day 4. See (58) for a full description of this experiment. **B)** Percent of untrimmed reads remaining after each sequential step of the metabarcoding pipeline for the same samples shown in (A). Forward and reverse reads are abbreviated as R1 and R2, respectively. **C)** Forward and reverse untrimmed read quality profiles for all samples. The mean and median quality scores for each base position are plotted as solid green and orange lines, respectively, and the 25% and 75% quality quartiles as dotted orange lines. The heatmap behind the line plots shows the basecall quality distribution for all reads in each dataset, with darker colors corresponding to more bases having that quality score.

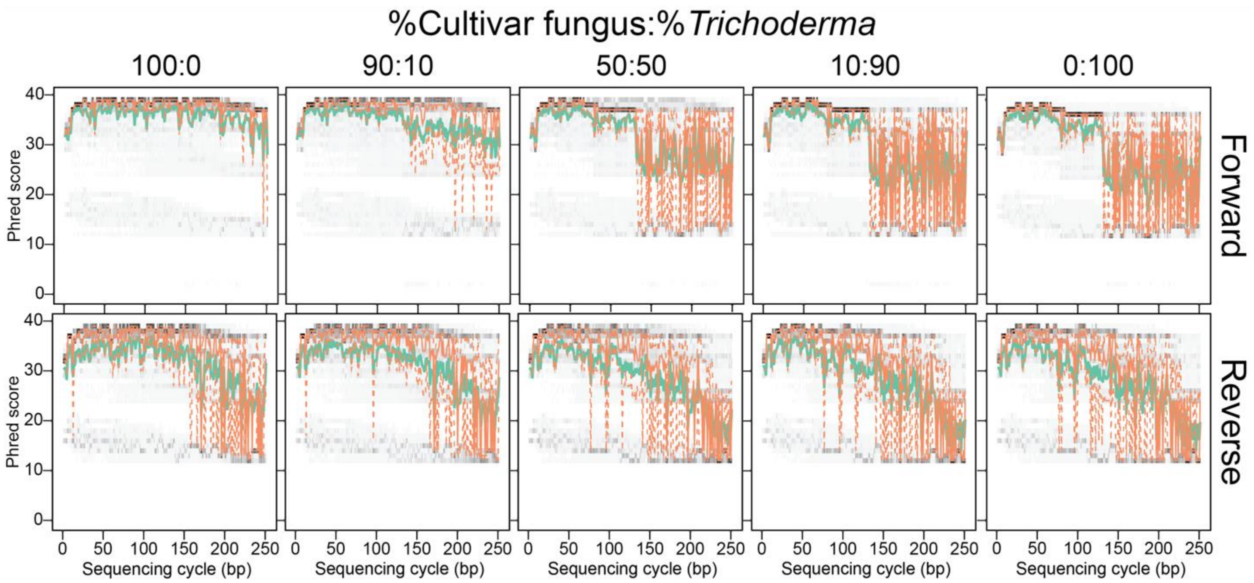


**<u>Figure S2</u>. Untrimmed forward (top) and reverse (bottom) read quality plots for the mock communities replicate the bias in *Trichoderma* read quality.** The mean and median quality scores for each base position are plotted as solid green and orange lines, respectively, and the 25% and 75% quality quartiles as dotted orange lines. The heatmap behind the line plots shows the basecall quality distribution for all reads in each dataset, with darker colors corresponding to more bases having that quality score.

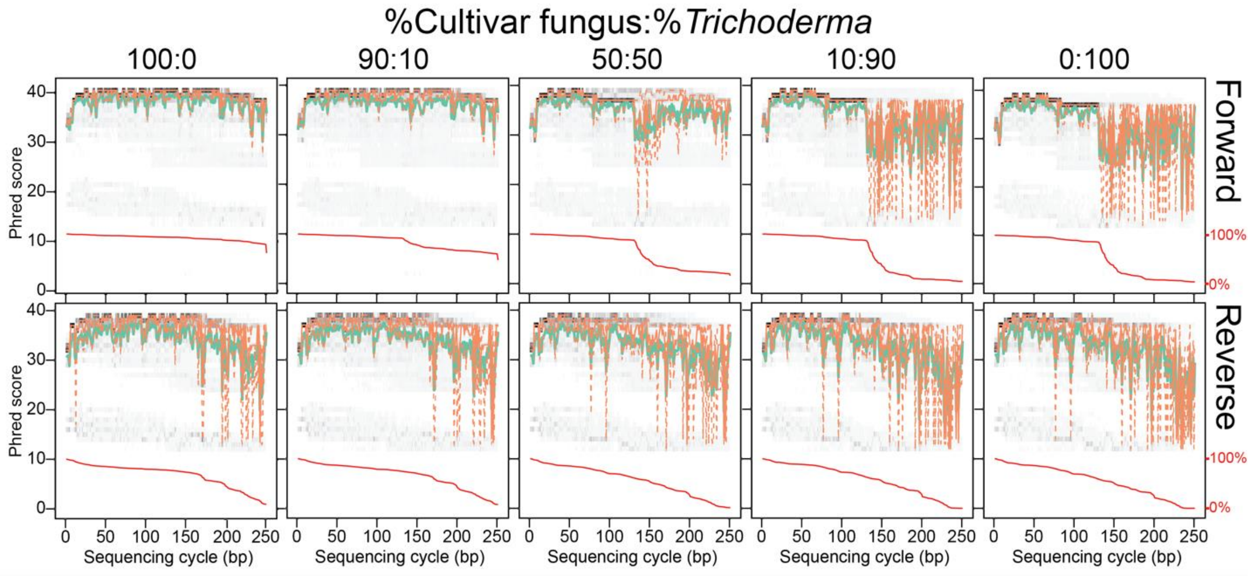


**<u>Figure S3</u>. Sliding window trimming improves the quality of forward (top) and reverse (bottom) reads in the mock community.** The mean and median quality scores for each base position are plotted as solid green and orange lines, respectively, and the 25% and 75% quality quartiles as dotted orange lines. The heatmap behind the line plots shows the basecall quality distribution for all reads in each dataset, with darker colors corresponding to more bases having that quality score. The red line below the quality scores indicates the percentage of reads of at least that sequence length that remain after trimming.

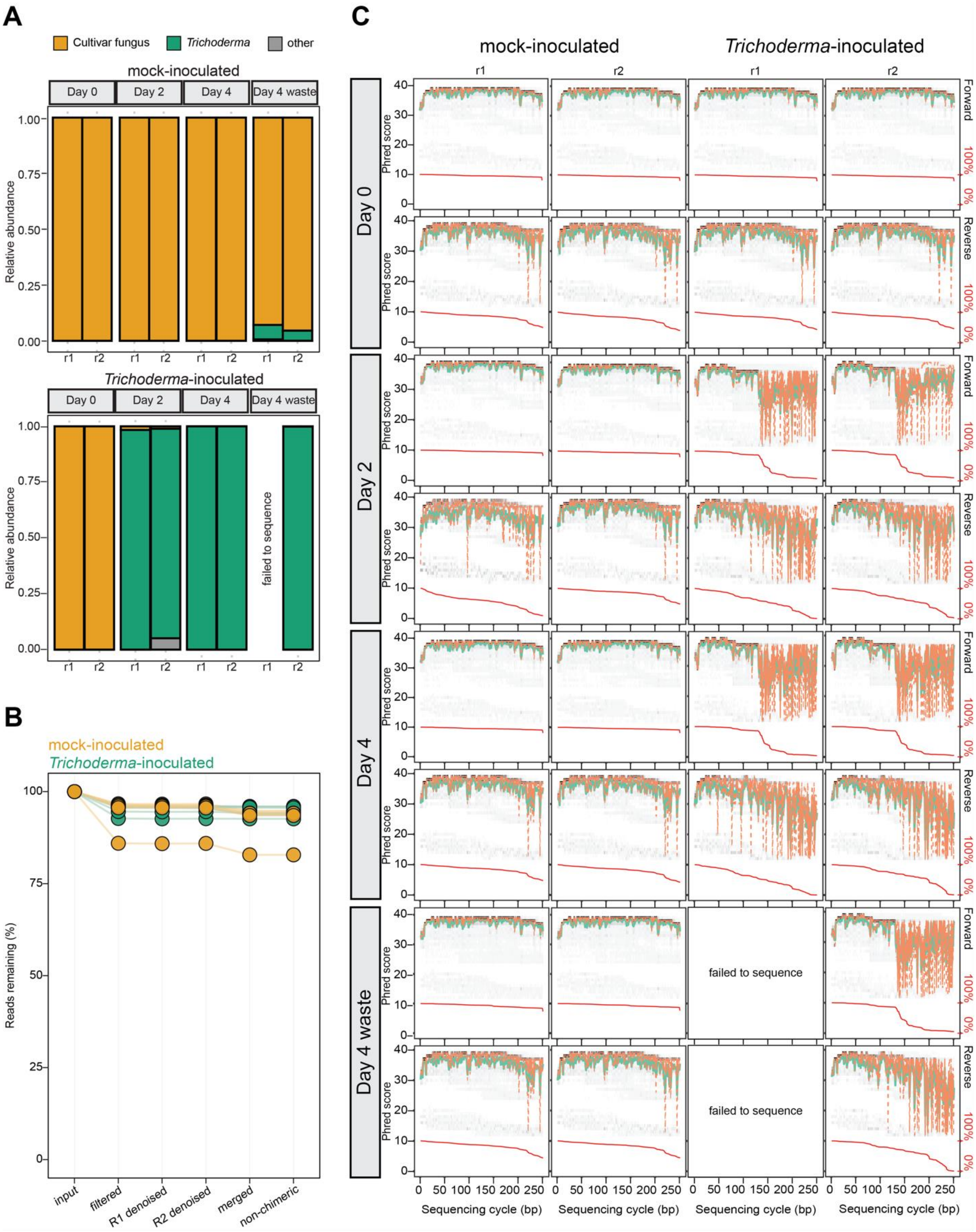
A
Cultivar fungus
Trichoderma
other
mock-inoculated
Day 0
Day 2
Day 4
Day 4 waste
Relative abundance
r1
r2
Trichoderma-inoculated
failed to sequence
B
mock-inoculated
Trichoderma-inoculated
Reads remaining (%)
input
filtered
R1 denoised
R2 denoised
merged
non-chimeric
C
mock-inoculated
Trichoderma-inoculated
Forward
Reverse
Phred score
100%
0%
failed to sequence
Sequencing cycle (bp)

**Figure S4. Sliding window trimming rescues *Trichoderma* ITS2 reads in the infection experiment. A)** Relative abundance stacked bar charts using trimmed reads for the time course *Trichoderma* infection experiment shown in Fig. 1 and Suppl. Fig. S1. Days 0, 2, and 4 follow inoculation with either *Trichoderma* spores (*"Trichoderma*-inoculated") or sterile PBS ("mock-inoculated"). Fungus garden waste (i.e., pieces removed from the fungus garden by the ants) was only collected on day 4. See (58) for a full description of this experiment. **B)** Percent of trimmed reads remaining after sequential steps of the metabarcoding pipeline for the same samples shown in A. Forward and reverse reads are abbreviated as R1 and R2, respectively. **C)** Forward and reverse read quality profiles for all samples. The mean and median quality scores for each base position are plotted as solid green and orange lines, respectively, and the 25% and 75% quality quartiles as dotted orange lines. The heatmap behind the line plots shows the basecall quality distribution for all reads in each dataset, with darker colors corresponding to more bases having that quality score. The red line below the quality score data shows the percentage of reads of at least that sequence length that remain after trimming.

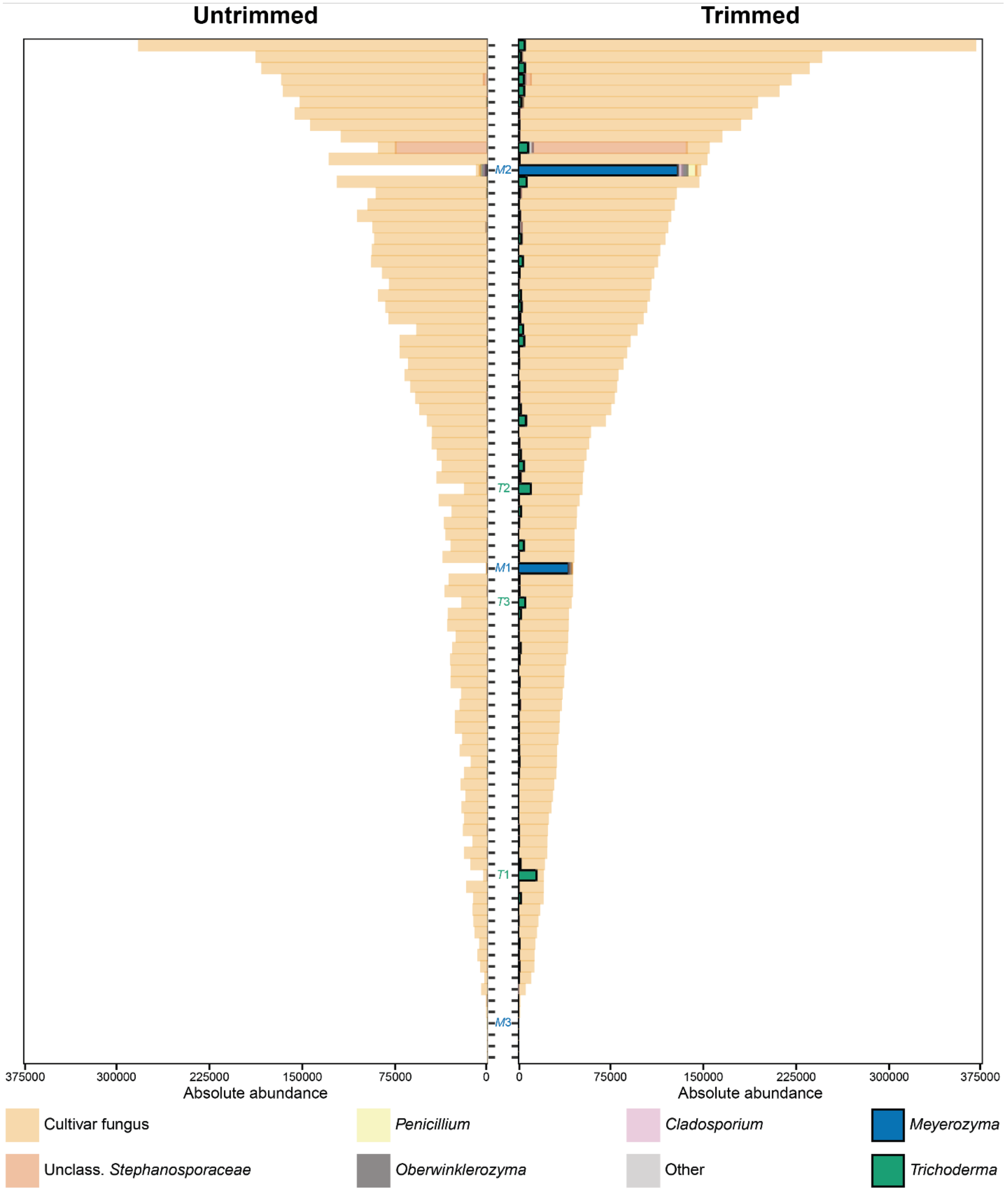

Untrimmed
Trimmed
M2
T2
M1
T3
T1
M3
375000
300000
225000
150000
75000
0
Absolute abundance
Cultivar fungus
Unclass. Stephanosporaceae
Penicillium
Oberwinklerozyma
Cladosporium
Other
Meyerozyma
Trichoderma

**Figure S5. Sliding window trimming increases the total number of reads that pass filtering and merging for all taxa in the environmental ITS2 dataset.** Absolute abundances of environmental ITS2 reads before (left) and after (right) trimming are ordered by decreasing absolute abundance of trimmed reads. The data used are the same as in Fig. 3, and *T*1-3 and *M*1-3 are the samples containing the three highest relative abundances of *Trichoderma* and *Meyerozyma*, respectively.

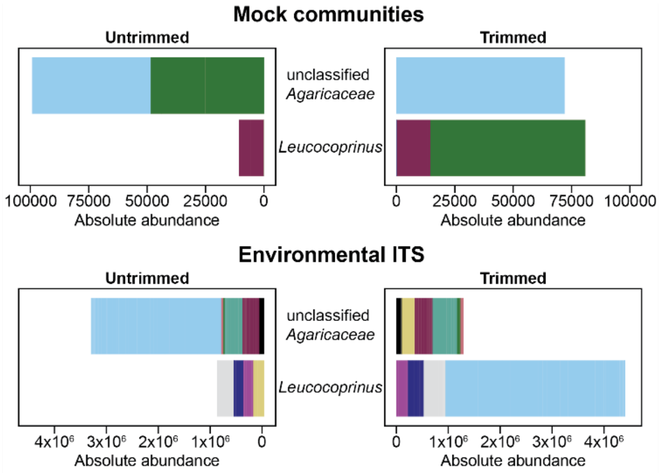


**Figure S6. Sliding window trimming (right) improves cultivar fungus classification in both the mock community and environmental datasets.** Individual ASVs are represented by different colors independently for each dataset. The mock community datasets (top) contained 5 cultivar fungus ASVs both before (left) and after trimming (right). The environmental dataset (bottom) contained 74 and 63 cultivar fungus ASVs before (left) and after trimming (right), respectively. Only the top 10 cultivar fungus ASVs of the trimmed environmental dataset are shown, comprising 92.5% of all cultivar fungus reads.

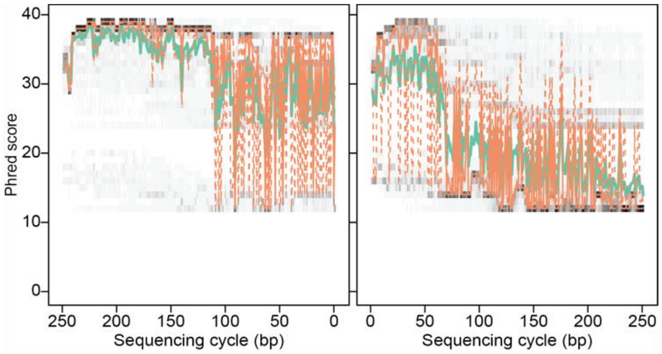


**<u>Figure S7</u>. *Clonostachys* has abnormally low read quality similar to, but distinct from, that of *Trichoderma* and *Meyerozyma*.** ITS2 read quality for genus *Clonostachys* forward and reverse reads are shown on the left and right, respectively. The mean and median quality scores for each base position are plotted as solid green and orange lines, respectively, and the 25% and 75% quality quartiles as dotted orange lines. The heatmap behind the line plots shows the basecall quality distribution for all reads in each dataset, with darker colors corresponding to more bases having that quality score.
